# Class-Controlled Copy-Paste Based Cell Segmentation for CoNIC Challenge

**DOI:** 10.1101/2022.03.02.482203

**Authors:** Heeyoung Ahn, Yiyu Hong

## Abstract

Muti-class cell segmentation in histopathology images is a challenging task. Here, we propose a copy-paste augmentation-based method for CoNIC challenge. As the challenge train data is severely class imbalanced. To deal with it, we copy all cell objects of train data and paste them to the train image on the fly while training model. The paste strategy is that we paste more cell objects of the insufficient classes and paste less cell objects for the sufficient classes. We experimented the method by stratified splitting train data in 4:1 ratio, the result shows the copy paste method can reach PQ 64.84 and mPQ 53.72, which improved and 0.66 compared to without copy pasted. Moreover, the improvements in those insufficient classes is more obvious.

## I. INTRODUCTION

This manuscript describes submitted method of team Arontier to CoNIC Challenge [1–2]. Our method mainly focus on how to deal with the class imbalance using copy-paste augmentation[1] with a little bit modification. We increase the number of cell objects for insufficient classes (Neutrophil, Plasma, Eosinophil) intensively and for those sufficient classes weakly using copy-paste augmentation, respectively. Importantly, considering there is no improvements when pasting cells to where they would be overlapped with other originally existed cells, we paste the cell objects to where they would not be overlapped.

## II. METHOD

### A. Extracting all cells and save

Unlike copy-paste augmentation[3] which do copy-paste in 2 different image pair, we want to paste cells from any images to target image. We extract all cell objects from all training set and saved them in a file in advance. As a result, we can efficiently use these cell objects and paste them on target train images when training our model on the fly.

### B. Source of pasted cells

For our copy-paste augmentation, we consider two method. One is source-level and the other is random-level. In source level, considering there are five image sources(i.e., consep, crag, dpath, glas, pannuke) in datasets, we paste cells from image of specific source to other target image of same source. For example, we extract cells from images of consep source and paste those cells to target images of same consep source. Unlike this, in random-level we extract cells from images of all sources and paste those cells to target images randomly without considering any sources. In our experiments, we found that random level yields better performance over source-level.

### C. Calculate candidate box region

For pasting cells to none-overlapping region, we firstly calculate the empty candidate box region to which the cells would be pasted. To this end, we stride a window in target image and calculate the total number of candidate box region to which cells would be pasted. Considering various size of cells, we select window size with 15-pixel width and height.

### D. Number of candidate box region to use and assigning different ratio for each class

After calculating the total number of candidate box region in each image, we set the ratio of how many boxes to use. We select random values of 0.1 interval between 0.1 and 1. Then we assign different ratio for each class according to the number of cells of a particular class in training images. Specifically, we assign different ratio of 0.3, 0.1, 0.05, 0.15, 0.3, 0.1 for each class because we want to deal with the insufficiency of particular classes. Finally if the pasted cells still overlap with other cells, we skip those cells.

### E. Augmentation for pasted cell

We experimented several augmentation method such as flip, scaling for cells which will be pasted. We found that scaling was not effective so we just use horizontal, vertical flip.

## III. TRAINING DETAILS

We follow the HoverNet[2] but with some differences. We replace the HoverNet[2] model structure with Unet++[3] and modify the structure so that model can output 3 outputs like HoverNet[2]. We select EfficientNet-b7[4] pretrained on ImageNet[5] for encoder of Unet++[3]. For Image Augmentation, we choose ColorJitter, Flip, Rotate, Blur, Gaussian Noise, Cutout [6], Cutmix[7]. We select Radam [10] optimizer with learning rate of 1e-3. During training, we downscale learning rate from 1e-3 to 1e-5 using step scheduler with 0.1 ratio every 100 epoch. For loss function, we choose same loss function that baseline of challenge [1] applied except for classification output. For classification output, we choose focal loss [11] with gamma 2, alpha 1. We train the model with batch size 8 for 300 epochs. Finally, we ensemble outputs of 5 models which were trained in 5-Fold cross validation setting.

## IV. RESULTS

For validating proposed copy-paste augmentation, we split datasets stratifiedly by 4:1 (train: valid) setting which is same as baseline of challenge. Table 1, 2 shows the results of applying proposed copy-paste augmentation for total classes and each class, respectively. When applied proposed copy-paste augmentation, for total classes PQ and mPQ improve by 0.6, 0.66 respectively and for each class PQ improve by 0.67, 0.32, 0.13, 1.08, 1.62, 0.14 respectively on validation set. Finally, Table 3 shows 5-Fold cross validation results of our model for submission.

**TABLE 1.** Results of model with / without proposed copy-paste for total classes on CoNIC challenge Dataset.

| Model | PQ | mPQ |
| --- | --- | --- |
| Unet++ | 64.24 | 53.06 |
| w/ Copy-Paste | 64.84( $\uparrow 0.6$) | 53.72( $\uparrow 0.66$) |

**TABLE 2.** Results of model with / without proposed copy-paste for each class on CoNIC challenge Dataset.

| Classes | Model | PQ |
| --- | --- | --- |
| 1. Neutrophil | Unet++ | 31.32 |
| | w/ Copy-Paste | 31.99( $\uparrow 0.67$) |
| 2. Epithelial | Unet++ | 63.49 |
| | w/ Copy-Paste | 63.81( $\uparrow 0.32$) |
| 3. Lymphocyte | Unet++ | 72.90 |
| | w/ Copy-Paste | 73.03( $\uparrow 0.13$) |
| 4. Plasma | Unet++ | 52.39 |
| | w/ Copy-Paste | 53.47( $\uparrow 1.08$) |
| 5. Eosinophil | Unet++ | 38.18 |
| | w/ Copy-Paste | 39.80( $\uparrow 1.62$) |
| 6. Connective | Unet++ | 60.04 |
| | w/ Copy-Paste | 60.18( $\uparrow 0.14$) |

**TABLE 3.** Results of model for submission.

| Fold | PQ | mPQ |
| --- | --- | --- |
| Fold1 | 64.91 | 53.49 |
| Fold2 | 65.35 | 54.01 |
| Fold3 | 65.65 | 54.45 |
| Fold4 | 65.09 | 54.98 |
| Fold5 | 65.77 | 53.95 |

**Fig1.**
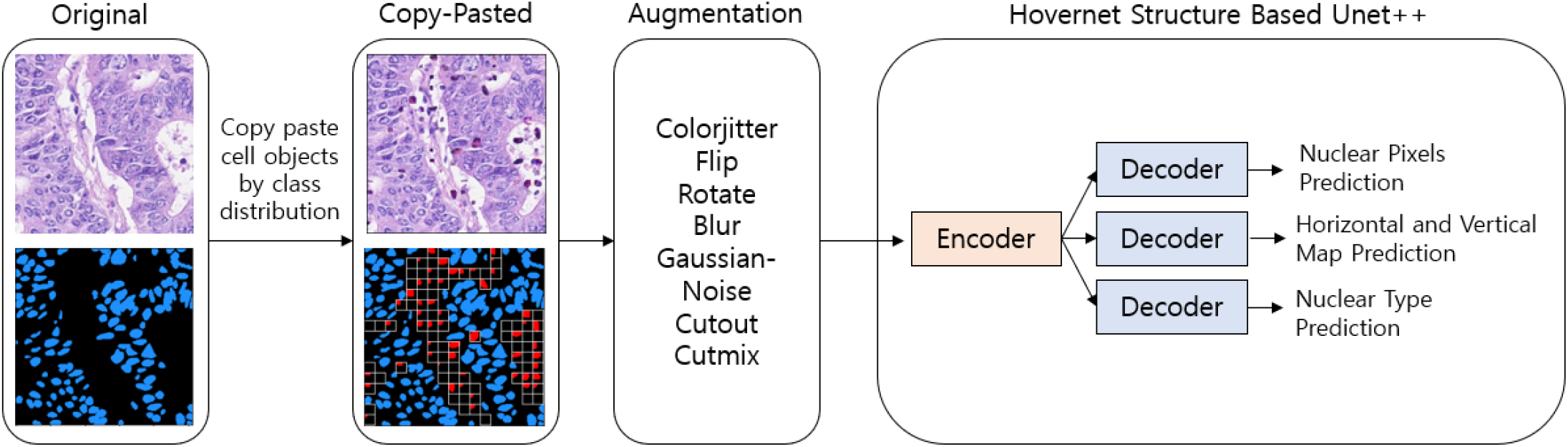
Overall Architecture of our proposed method.

